# Expression of MMP-11, an estrogen-suppressed gene in MCF-7 cells, is elevated upon acquisition of tamoxifen resistance

**DOI:** 10.1101/2025.02.13.638037

**Authors:** Ningthoujam Sonia, Sreeja Mondal, Anil Mukund Limaye

**Author notes:** Corresponding author Anil Mukund Limaye, Department of Biosciences and Bioengineering, Indian Institute of Technology Guwahati, Guwahati 781039, Assam, India. **Authors’ email** Ningthoujam Sonia, Sreeja Mondal. **Author Contributions:** NS: cell culture experiments, data acquisition, preparation of manuscript; SM: RNAseq data analysis, AML: Funding acquisition, manuscript editing, conceptualization and overall supervision. **Competing Interest Statement:** The authors declare no competing interests.

## Abstract

Matrix metalloproteinase-11 (MMP-11), also known as stromelysin-3, is a member of a large family of zinc-dependent proteinases. Known to be expressed in breast tumor stroma, the epithelium, and the associated mononuclear inflammatory cells, its expression is associated with invasion, metastasis, and poor clinical outcomes and survival rates. Despite the known effects of estrogen on extracellular matrix remodeling, and breast cancer metastasis, the role of MMP-11 therein is not understood. Here, we show that estrogen suppresses MMP-11 expression in MCF-7 breast cancer cells via a mechanism that involves binding of the estrogen receptor α (ERα) to an estrogen response element located in the intron-1 of MMP-11. ER_α_ knockdown alone upregulates MMP-11 expression, highlighting its crucial role in this regulatory axis. Additionally, the anti-estrogen tamoxifen (4-OHT) blocks estrogen-mediated MMP-11 suppression. However, prolonged 4-OHT exposure or 4-OHT resistance leads to increased MMP-11 expression. This suggests that MMP-11 may play a role in 4-OHT resistance, potentially leading to a more aggressive tumor phenotype. Our study underscores the importance of estrogen-ER_α_ signaling in breast cancer progression, emphasizing the need for further investigation into MMP-11’s role in 4-OHT resistance and cancer metastasis.

## 1. Introduction

Matrix metalloproteinase-11 (MMP-11), also known as stromelysin-3 (ST-3), is a clinico-pathologically significant member of a large family of extracellular matrix (ECM)-remodeling and zinc-dependent endopeptidases, called matrix metalloproteinases (MMPs). Initially identified in tumor-adjacent stroma [1], MMP-11 is also expressed in tumor cells [2–4]. Unlike most other members of the MMP family, it is activated by furin-like convertases, and spares conventional MMP substrates like elastin, laminin, and fibronectin [1]. However, it cleaves α3 chain of collagen VI, insulin-like growth factor binding protein-1 (IGFBP-1), and serine protease inhibitor _α_1-antitrypsin [1].

MMP-11 is implicated in the progression of breast, prostate, and lung cancers [1]. It is frequently overexpressed in breast tumors [3], and is associated with proliferation and malignant progression [5–7]. High expression of MMP-11 in breast tumors and stromal tissues is correlated with poor prognosis, characterized by reduced overall, and relapse-free survival [1,8]. Moreover, its expression is associated with various clinicopathological parameters, including ER [2] and PR status, tumor stage, nodal metastasis [4], and prognostic markers like Ki-67 and Bcl-2 [3]. Additionally, MMP-11 expression in mononuclear inflammatory cells (MICs) significantly predicts clinical outcome; correlating with reduced relapse-free survival, and overall survival rates. MMP-11 expression in MICs may be associated with pre-invasive changes in breast cancers. Tumors infiltrated by MMP-11 positive MICs express elevated levels of IL-1, IL-5, IL-6, IL-17, IFNβ, and NFκB [9].

MMPs, in general, by virtue of their ability to remodel the extracellular matrix, promote invasion and metastasis. Paradoxically, MMP-11 appears to positively or negatively impact tumorigenesis and tumor progression (reviewed by [1]). Experiments on subcutaneous tumor models in nude mice have revealed tumor-promoting effects of MMP-11 in NIH3T3 and MCF-7 cells. However, the rate of proliferation of MMP-11 overexpressing MCF-7 tumors was no different from those produced by parental MCF-7 cells [10]. Consistent with the tumor-promoting role, MMP-11 deficient MMTV-ras transgenic mice develop fewer and smaller primary tumors. However, these mice have elevated lung metastatic lesions; demonstrating the dual role of MMP-11 in breast cancer [11].

MMP-11 is regulated at multiple levels; from transcriptional regulation to pericellular inhibition [1,12,13]. In the context of breast cancer, PKD1 [14], Gli-1 [1], and ECM-associated growth factors, such as IGF-1 [15], regulate MMP-11. Total estrogen exposure is a major factor in the etiology of breast cancer. Estrogen not only promotes the proliferation of breast cancer cells, but also modulates the tumor microenvironment (TME) that is associated with tumor progression [16,17]. Several MMPs are regulated by estrogen [18,19]. Despite the recognized prognostic significance of MMP-11 expression in breast cancer, its relationship with the estrogen-ER_α_ signaling axis remains unexplored.

Here we show that the natural estrogen, 17β-estradiol (E2), down modulates the expression of MMP-11 expression in ERα-positive and E2-responsive MCF-7 breast cancer cells via a mechanism that involves ERα. 4-hydroxy tamoxifen (4-OHT), a selective estrogen receptor modulator, inhibits estrogen-mediated suppression of MMP-11 in MCF-7 cells. However, 4-OHT resistance, or prolonged exposure of MCF-7 cells to 4-OHT, is associated with increased MMP-11 expression.

## 2. Materials and methods

### 2.1. Plasticware, chemicals, and reagents

Cell culture plasticware was sourced from Eppendorf (Hamburg, Germany). Dulbecco’s Modified Eagle’s Medium (DMEM) with and without phenol red, Dulbecco’s phosphate-buffered saline (DPBS), fetal bovine serum (FBS), charcoal-stripped fetal bovine serum (csFBS), trypsin-EDTA, and antibiotic solution were purchased from HiMedia Laboratories (Mumbai, India). 17β-estradiol (E2, Cat. No. E8875), 4-OHT (Cat. No. H7904), and propyl pyrazole triol (PPT, Cat No. H6036) were from Sigma Aldrich (MO, USA).

### 2.2. Cell line and cell culture

MCF-7 breast cancer cells, sourced from the National Center for Cell Science, Pune, India, were cultured in phenol red-containing DMEM, which was supplemented with 10% (v/v) heat-inactivated FBS, 100 U/ml penicillin, and 100 μg/ml streptomycin (M1 medium). The cells were maintained at 37°C under a humidified environment with 5% CO_2_. For treatment with estrogenic ligands, cells were supplied with phenol red-free DMEM, which was supplemented with 10% (v/v) heat-inactivated csFBS, 100 U/ml penicillin, and 100 μg/ml streptomycin (M2 medium).

### 2.3. Experimental design

#### 2.3.1. Time-course experiment

2×10^5^ MCF-7 cells were grown in 35 mm dishes with M1 until 60-70% confluent. Cells were washed with DPBS, and fed with M2. After 24 h, cells were then treated with ethanol (vehicle) or 10 nM E2 in M2 for varying periods of time (0 to 96 h), with replacement of the treatment medium every 48 h.

#### 2.3.2. Dose-response experiment

2×10^5^ MCF-7 cells were grown in 35 mm dishes using M1 until 60-70% confluent. Following a DPBS wash, they were fed with M2 for 24 h. Thereafter, cells were treated with ethanol (vehicle) or varying concentrations of E2 (0.1 to 100 nM) in M2 for 24 h.

#### 2.3.3. Effect of 4-OHT

2×10^5^ MCF-7 cells were grown in 35 mm dishes for 24 h in M1. Cells were washed with DPBS, and fed with M2 for 24 h. Thereafter, the cells were treated with ethanol (vehicle), 10 nM E2, or 10 nM PPT in M2, alone or in combination with 1 μM 4-OHT for 24 h.

#### 2.3.4. Effect of ER_α_ knockdown

MCF-7 cells were grown in 6-well plates (2×10^5^ cells/well) using M1 for 24 h. The cells were transfected with ER_α_-specific siRNA (Cat No. 4392420, Thermo Scientific, PA, USA) or scrambled siRNA (Cat No. AM4611, Thermo Scientific, PA, USA) for 24 h using Lipofectamine RNAiMax (Cat No. 13778075, Thermo Scientific, PA, USA), as per manufacturer’s instructions. The cells were washed with DPBS, and fed with M2 for 24 h. Thereafter, the cells were treated with ethanol (vehicle) or 10 nM E2 in M2 for 24 h.

### 2.4. RNA Isolation, cDNA synthesis and RT-qPCR

Total RNA was extracted using a reagent following the method of Chomczynski and Sacchi [20], and its concentration was determined using BioSpectrometer basic (Eppendorf, Hamburg, Germany). 2 μg of total RNA was reverse transcribed, and cDNA equivalent to 20 ng of total RNA served as a template for amplification with gene-specific primers (Supplementary Table S1) in AriaMx Real-time PCR System (Agilent, CA, US). The expression levels of each gene were analyzed using the ΔΔCt method [21], with RPL35a as the internal control.

### 2.5. Western blotting

Total protein was extracted from cells using RIPA buffer. Alternatively, total protein was extracted from the organic phase that is separated during total RNA isolation as described by Likhite et al. [22], with slight modifications. Total protein was quantified by Lowry s method [23]. 30 μg of protein samples were resolved in 10% denaturing SDS-PAGE, transferred to 0.45 µ nitrocellulose membrane (Cat. No. SF110B, HiMedia Laboratories, Mumbai, India), and blocked with either tris-buffered saline containing 2% Tween 20 (TBST) or 1% (w/v) gelatin in tris-buffered saline containing 0.05% v/v Tween 20 (TBST) for 1 h at room temperature. The blots were then probed with polyclonal anti-MMP-11 antibody (Supplementary data 1) for 2 h or anti-H3 antibody (Cat. No. BB-AB0055, BioBharati LifeScience, India) for 1h at room temperature. The blots were washed with TBST (6 × 5 min). Blots were then probed with goat anti-rabbit HRP conjugated secondary antibody (Cat. No. 7074S, Cell Signaling Technology, MA, USA) for 1h, washed with TBST (6 × 5 min), and developed using Clarity Western ECL Substrate (Bio-Rad, California, US). Images were captured with the ChemiDoc XRS+ system (Bio-Rad, California, US).

### 2.6. ChIP-seq analysis

The ER_α_ occupancy within the MMP-11 gene was analyzed using publicly accessible ChIP-seq data from the Sequence Read Archive (SRA) as described earlier [24]. For this study, we selected the ChIP-seq data (SRA accession ID: ERP000380), which were generated from chromatin samples of MCF-7 cells treated with E2 (ID: ERR022026), or vehicle (ID: ERR022025), followed by immunoprecipitation with anti-ER_α_ antibody. Analysis was carried out using Galaxy, a web-based platform (https://usegalaxy.org/) [25]. The ER_α_ occupancy was then viewed with the UCSC genome browser.

### 2.7. Chromatin immunoprecipitation (ChIP)

Chromatin immunoprecipitation was carried out as described earlier [24]. 80 µg of chromatin samples were pre-cleared using Protein G plus-Agarose beads (Cat No. IP04-1.5ML, Merck Millipore, USA) pre-coated with bovine serum albumin (BSA) and herring sperm DNA for 2 h. A fraction (5%) of the pre-cleared chromatin samples were set aside as input. The remaining portions were incubated with ER_α_-specific antibody (Cat. No.8644S, Cell Signaling Technology, MA, USA) or rabbit IgG antibody (BioBharati LifeScience, India) at 4°C, for 4 h. The immunoprecipitates were washed with a series of wash buffers, and reverse-crosslinked as described earlier [24]. Immunoprecipitated DNA was purified using column, and ER_α_ occupancy was evaluated by PCR using primers (Supplementary Table S1) specific to the estrogen-response elements within pS2 (positive control) or MMP-11 locus.

### 2.8. RNA-seq data and differential gene expression analysis

Raw RNA-seq data (GSE164529) corresponding to parental, and 4-OHT-resistant MCF-7 cells, which were generated by Jones CJ et al. [26], was downloaded. The read quality was assessed and processed to remove adapters, and filtered to retain good-quality reads as described earlier [27]. The trimming parameters included-ILLUMINACLIP:TruSeq3-PE.fa:2:30:10:2:keepBothReads, HEADCROP:15, LEADING:3, TRAILING:3, SLIDINGWINDOW:4:30, MINLEN:30. Reads were aligned to the human genome, and gene expression data, in the form of read counts, were obtained and processed as previously described [27]. The read-count data were examined through PCA plots and correlation heatmaps. The Wald statistic was used to identify differentially expressed genes using the threshold values of 0, and 0.05 for log_2_FoldChange, and adjusted p-value (padj), respectively.

### 2.9. Long-term 4-OHT treatment of MCF-7 cells

MCF-7 cells were seeded at a density of 2×10^6^ cells per 100 mm dish and grown in M1 medium. The cells were treated chronically with 1 μM 4-OHT for 26 days. Ethanol (vehicle)-treated cells served as control.

### 2.10. Statistical analysis

To identify differentially regulated genes in 4-OHT-resistant cells, we employed DESeq2, with Wald statistic, with _α_ equal to 0.05. Subsequently, we applied FDR correction with a 5% cut-off. For two group data, we used Welch’s two-sample t-test. ER_α_ knockdown data underwent two-way ANOVA analysis after confirming homogeneity of variance via the Levene test, and assessing normality with the Shapiro-Wilk test. One-way ANOVA was utilized for other multiple-group data. All statistical tests were performed at a 5 % level of significance.

## 3. Results

### 3.1. Estrogen regulates MMP-11 expression

Fig. 1 shows the results of time-course, and dose-response studies that examined the effect of E2 on MMP-11 expression in MCF-7 breast cancer cells. At all time points tested (6 to 96 h), 10 nM E2-treated cells expressed significantly lower levels of MMP-11 mRNA (Fig. 1a). All the tested concentrations of E2 (0.1 to 100 nM) significantly reduced MMP-11 mRNA (Fig. 1c). The reduction in MMP-11 protein expression paralleled the decrease in MMP-11 mRNA in both time-course (Fig. 1b), and dose-response experiments (Fig. 1d).

**Fig. 1.**
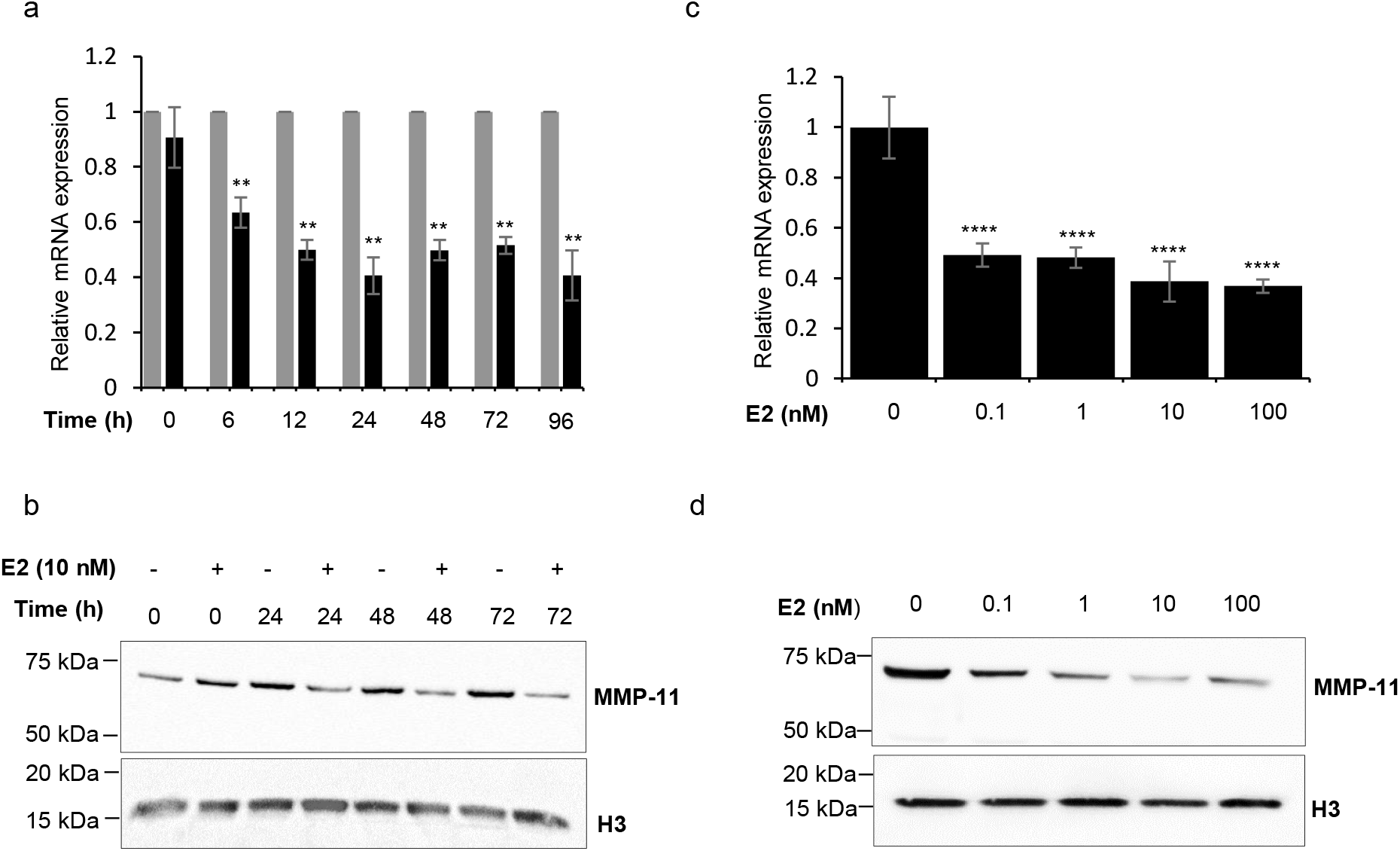
E2 regulates MMP-11. **a,b**. Time-course experiment. MCF-7 cells were treated with 10 nM E2 or vehicle (0.1 % EtOH) for the indicated period of time. Total RNA was isolated and subjected to RT-qPCR using MMP-11-specific primers (a). RPL35a served as an internal control. For each time point, the normalized MMP-11 expression in control samples was set to 1, and those in treated samples were expressed relative to control. Bars represent mean relative expression ± sd (n=3 biological replicates). The data for each time point were analyzed using Welch’s two-sample *t*-test. **denotes significant result relative to control (*p*<0.01). Western blot analysis of MMP-11 protein expression (b). Total protein was extracted, and 30 µg of each protein sample was subjected to western blot using MMP-11 specific antibody. H3 served as the internal control. **c,d**. Dose-response experiments. MCF-7 cells were treated with vehicle (0.1 % EtOH) or indicated concentrations of E2 for 24 h. Total RNA was isolated and subjected to RT-qPCR using MMP-11-specific primers (c). RPL35a served as an internal control. The expression level of a control sample was set to one, and MMP-11 expression in each sample was expressed relative to this control. Bars represent mean relative expression ± sd (n=3 biological replicates). The data were analyzed using ANOVA followed by Tukey’s HSD. ****denotes significant results relative to control (*p*<0.0001). Western blot analysis of MMP-11 protein expression (d). Total protein was extracted, and 30 µg of each protein sample was subjected to western blot using MMP-11 specific antibody. H3 served as the internal control. The chemiluminescence images show the expression levels of MMP-11 in samples from one out of 3 biological replicate experiments.

### 3.2. Estrogen regulates MMP-11 expression via ER_α_

To examine whether the E2-regulation of MMP-11 involves ER_α_, MCF-7 cells were treated with vehicle, E2, or PPT (an ERα agonist), alone or in combination with 4-OHT, for 24 h. Both E2 and PPT independently suppressed MMP-11 mRNA expression (Fig. 2a, bar 1 vs bars 2 and 3). While 4-OHT marginally but significantly reduced MMP-11 mRNA expression, it significantly blocked the E2- or PPT-mediated suppression of MMP-11 mRNA. E2 alone reduced MMP-11 mRNA expression by 66%, whereas, in the presence of 4-OHT, it reduced the expression by 38% (Fig. 2a, bar 1 vs bars 2 and 5). Similarly, PPT alone reduced the MMP-11 mRNA expression by 73.8%, whereas in the presence of 4-OHT, it reduced the expression by 23% (Fig. 2a, bar 1 vs bars 3 and 6). Western blots revealed that 4-OHT also blocked E2- or PPT-mediated suppression of MMP-11 protein (Fig. 2b).

**Fig. 2.**
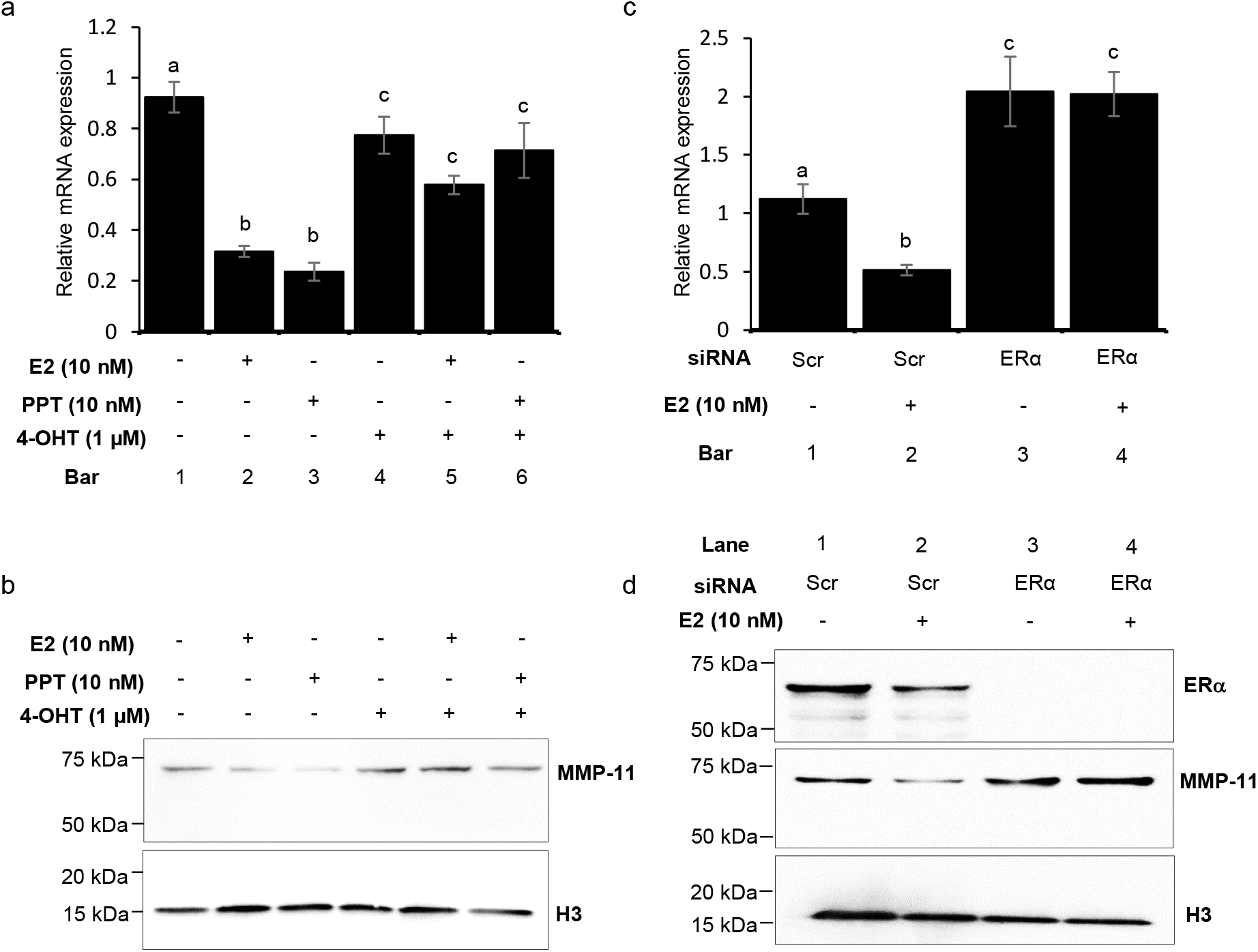
Estrogen regulation of MMP-11 is mediated via ERα. **a,b**. Effect of 4-OHT on E2- or PPT-mediated expression of MMP-11. MCF-7 cells were treated with 10 nM E2 or 10 nM PPT, alone or in combination with 1 μM 4-OHT. The experiment also included cells treated with only 1 μM 4-OHT. Total RNA was isolated and subjected to RT-qPCR using MMP-11-specific primers (a). RPL35a served as an internal control. The expression level of a control sample was set to one, and MMP-11 expression in other samples was expressed relative to this control. Bars represent mean relative expression ± sd (n=3 biological replicates). The data were analyzed using ANOVA followed by Tukey’s HSD. Western blot analysis of MMP-11 protein expression (b). Total protein was extracted, and 30 µg of each protein sample was subjected to western blot using MMP-11 specific antibody. H3 served as the internal control. **c,d**. Effect of ERα knockdown on E2 regulation of MMP-11. MCF-7 cells were transfected with scrambled (Scr) or ERα-specific siRNA and incubated for 24 h, followed by treatment with vehicle (0.1 % EtOH) or 10 nM E2 for 24 h. Total RNA was isolated and subjected to RT-qPCR using MMP-11-specific primers (c). RPL35a served as an internal control. The expression level of a control sample was set to one, and MMP-11 expression in each sample was expressed relative to this control. Bars represent mean relative expression ± sd (n=3 biological replicates). The data were analyzed using two-way ANOVA to determine the main effects of estrogen, ERα siRNA, or their interaction, followed by Tukey’s HSD post-hoc test. Western blot analysis of MMP-11 protein expression (d). Total protein was extracted, and 30 µg of each protein sample was subjected to western blot using MMP-11 or ERα-specific antibodies. H3 served as the internal control. Letter codes (a, b, c, and d) above the bars denote statistical differences between treatment pairs. The chemiluminescence images show the expression levels of MMP-11 in samples from one out of 3 biological replicate experiments

We also tested the effect of siRNA-mediated knockdown of ERα on MMP-11 expression. Its knockdown alone, reflected by significantly reduced expression of ERα protein (Fig.2d, lane 1 vs lane 3), significantly increased the expression of MMP-11 mRNA (Fig. 2c, bar 1 vs bar 3). However, ERα knockdown completely inhibited E2-mediated suppression of MMP-11 mRNA (Fig. 2c, bar 1, bar 3 and 4). Western blotting data (Fig. 2d) are consistent with these results.

### 3.3. Estrogen treatment enhances ER_α_ binding at the MMP-11 locus

ERα is a ligand-dependent transcription factor, that interacts with estrogen response elements (EREs) in the target-gene regulatory region. Through three independent approaches, we examined whether E2-mediated suppression of MMP-11 mRNA was associated with increased ERα occupancy in the regulatory regions of MMP-11. We scanned the MMP-11 locus with the JASPAR transcription factor binding site prediction tool, to predict a potential ERE in intron-1 of MMP-11 (Fig. 3a). We then analyzed ChIP-seq data obtained from control and E2-treated MCF-7 cells. Enriched ERα binding to the intron-1 of the MMP-11 gene was found in E2-treated cells, but not in the control cells (Fig. 3a, indicated by the red rectangle). This ERα-enriched region overlapped with the ERE-containing sequence predicted by JASPAR. These findings were independently validated by ChIP experiments employing ERα-specific antibodies and PCR primers designed specifically to amplify a fragment of 140 bps within the ERα-enriched region revealed by ChIP-seq data (Fig. 3b).

**Fig. 3.**
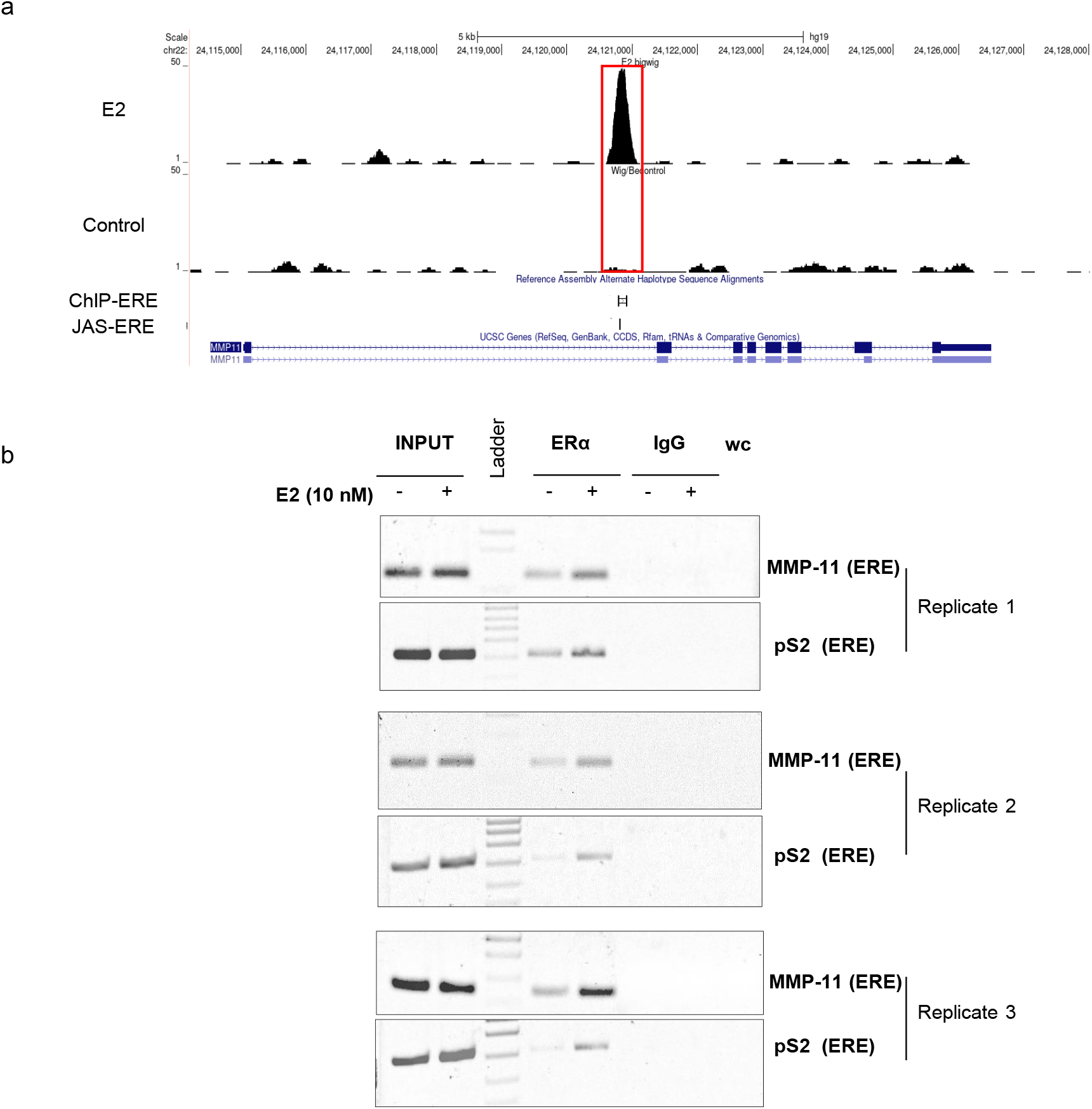
E2 enhances ER_α_ occupancy at the MMP-11 locus. **a**. ChIP-seq data from MCF-7 cells treated with vehicle (0.1 % EtOH) or 10 nM E2 (SRA accession no. ERP000380) were analyzed using Galaxy. The ERα binding across the MMP-11 gene were visualized in the UCSC genome browser. The major peak of ERα binding shown in red colored rectangle is observed only in the estrogen-treated sample. The presence and location of an ERE predicted by JASPAR is indicated in the JAS-ERE track. **b**. Validation of ERα binding using ChIP assay. Sonicated chromatin samples from MCF-7 cells treated with vehicle (0.1 % EtOH) or 10 nM E2 were immunoprecipitated with ERα-specific or normal IgG antibody (negative control). Immunoprecipitated DNA was subjected to PCR using primers specific to the MMP-11 (ChIP-ERE).

### 3.4. 4-OHT resistance of MCF-7 cells is associated with enhanced MMP-11 expression

Jones CJ et al [26], generated endocrine-resistant breast cancer cells from the parental MCF-7 cells by chronic exposure to endoxifen, 4-OHT, and fulvestrant (ICI182780). They characterized the transcriptomic profiles of these resistant cells, and vehicle-treated controls using the RNA-seq approach. Their RNA-seq data were accessed through the Gene Expression Omnibus using the accession number GSE164529. This dataset provided an opportunity to examine and compare the expression of MMPs in 4-OHT-resistant and vehicle-treated MCF-7 cells. We reanalyzed the RNA-seq data as described in section 2.8. Differentially expressed genes were identified by analyzing the raw count data using DESeq2. Applying Wald statistics with 5 % FDR (padj < 0.05) and log_2_FC cut-off of +/-1, 2963 genes (Fig. 4a) were found to be differentially expressed between 4-OHT-resistant and vehicle-treated MCF-7 cells. Among these were MMP-9, MMP-11, MMP-16, and MMP-17. The normalized counts for these MMPs in each of the biological replicates of vehicle-treated and 4-OHT-resistant cells are shown in Fig. 4b, which shows that MMP-9 and MMP-11 mRNA expression levels were higher in 4-OHT-resistant cells. On the other hand, the mRNA expression levels of MMP-16, and MMP-17 were lower. To independently validate the observation on MMP-11 expression, MCF-7 cells were subjected to chronic 4-OHT treatment for 26 days. The expression level of MMP-11 mRNA was then compared to that in MCF-7 cells treated with vehicle. As shown in Fig. 4c, the expression of MMP-11 mRNA was significantly higher in 4-OHT-treated MCF-7 cells, compared to those treated with vehicle. The MMP-11 protein expression was also higher in 4-OHT-treated cells (Fig. 4d).

**Fig. 4.**
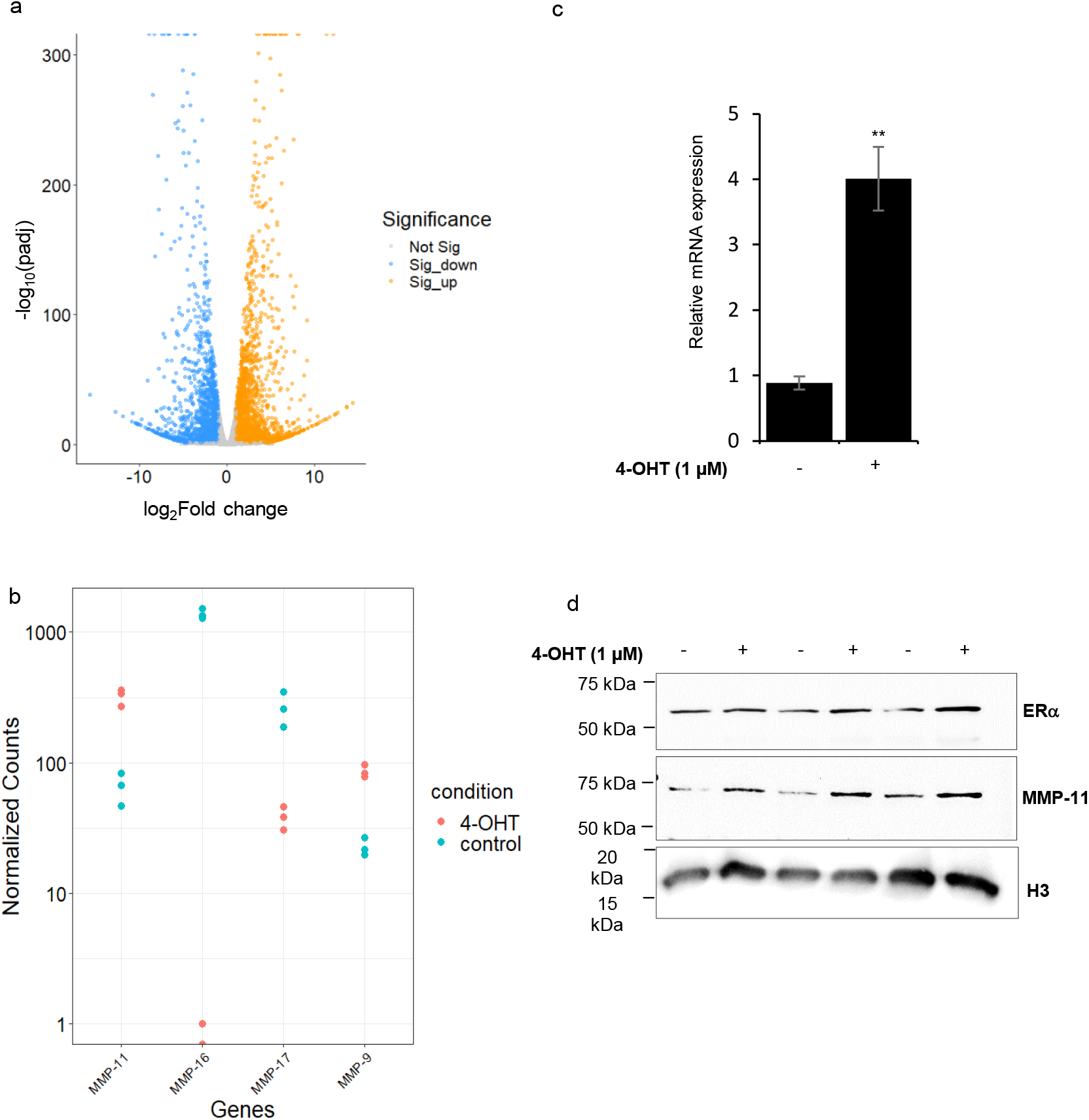
MMP-11 expression is enhanced in OHT-resistant cells. **a,b**. Reanalysis of the RNA-seq data (GSE164529). Volcano plot (a), showing significantly 1723 upregulated (orange dots, log2FC >1) genes, and 1240 downregulated genes (blue dots, log2FC < −1) in 4-OHT-resistant cells. Normalized counts of the indicated MMPs (b) in each of the replicate samples (n = 3). **c,d**. Effect of chronic treatment of MCF-7 cells with 4-OHT on MMP-11 expression. Total RNA was isolated and subjected to RT-qPCR using MMP-11-specific primers (c). RPL35a served as an internal control. The expression level of a control sample was set to one, and MMP-11 expression in other samples was expressed relative to this control. Bars represent mean relative expression ± sd (n=3 biological replicates). The data were analyzed by Welch’s two-sample t-test. **denotes significant result (p<0.01) compared to control. Western blot analysis of MMP-11 protein expression (d). Total protein was extracted, and 30 µg of each protein sample was subjected to western blot using MMP-11 or ERα-specific antibodies. H3 served as the internal control. The chemiluminescence images show the expression levels of MMP-11 in samples from one out of 3 biological replicate experiments.

## 4. Discussion

The estrogen-ER_α_ signaling axis is crucial for the growth and proliferation of breast epithelial cells, and plays a key role in breast tumorigenesis and metastasis. Evidence suggests that ER_α_ is a key mediator in the interaction between cancer cells and the TME, contributing to ECM remodeling and metastasis [16,17]. MMPs are critical for ECM degradation, and the promotion of tumor invasion. Aberrant expression of MMPs in breast cancer is associated with poor prognosis [8]. Several researchers have explored estrogen regulation of MMP-1, MMP-9, and MT1-MMP in breast cancer, emphasizing the involvement of E2-ER_α_ signaling [18,19]. The present study demonstrates that estrogen suppresses the expression of MMP-11 in MCF-7 cells. The time- and dose-dependent regulation of MMP-11 by estrogen or PPT, as well as the abrogation of estrogen-mediated effects by ER_α_-specific siRNA, indicates the role of ER_α_. ChIP experiments further revealed increased ER_α_ binding to the regulatory elements of MMP-11, indicating estrogen’s role in suppressing MMP-11 transcription. Reduced MMP-11 mRNA was also reflected in reduced MMP-11 protein. Therefore, it is likely that the estrogenic control of the TME during breast cancer progression is mediated at least in part via MMP-11.

We also performed similar experiments in T47D cells to examine whether E2-mediated modulation of MMP-11 is universal. Unlike MCF-7 cells, no change in MMP-11 expression was observed in T47D cells at early time points (6, 12, and 24 h). However, prolonged E2 treatment significantly increased MMP-11 mRNA levels at 48, 72, and 96 h (Supplementary figure 1). This variation in response to E2 was also observed in our previous study [24], where estrogen differentially regulated cystatin A in these two cell lines. Although the precise reason for the differences in MMP-11 regulation remains unclear, it may be attributed to the phenotypic differences resulting from the distinct gene expression profiles between the two cell lines [28]. Further investigation is needed to determine the mechanisms driving this variation. Despite these differences, ER_α_ depletion upregulated MMP-11 in both MCF-7 (Fig.2c) and T47D cells (Supplementary figure 1), suggesting that ERα tightly controls the basal, steady-state levels of MMP-11 expression in ER+ breast cancer cells. Clinically, this is relevant, as high MMP-11 expression in breast tumor cells is associated with ER-negative status [29], poor prognosis [2,3,5], and increases tumor aggressiveness [4].

MMP-11 facilitates metastasis by promoting cell invasion, and migration [5], and is highly expressed in metaplastic carcinomas, which are characterized by epithelial-to-mesenchymal transition (EMT) [30]. Furthermore, MMP-11 in breast cancer cells, via its paracrine signaling, stimulates MMP-11 expression in adjacent fibroblast cells [15], potentially fostering a tumor-promoting microenvironment. Bouris et al [18] also reported that ERα depletion induces the expression of MT1-MMP, MMP-1, MMP-2, and MMP-7, resulting in EMT phenotypes. Thus, MMP-11 upregulation upon ERα depletion may contribute to tumor aggressiveness, and EMT. This notion is further supported by observations that anti-estrogens like ICI 164,384 or ICI 182,780 increase cell invasiveness by inhibiting the protective effects of estrogen and ERα, likely by reducing ERα expression [31]. Consequently, antiestrogen treatments might inadvertently induce metastasis by suppressing ERα expression and altering MMPs expression.

While MMP-11’s role in promoting invasiveness and metastasis is well-established, its involvement in endocrine resistance remains poorly understood. The emerging link between metastasis and endocrine resistance, due to their shared molecular pathways [32], prompted us to investigate MMP-11’s role in 4-OHT resistance. Re-analysis of RNA-seq data (GSE164529) in this study revealed elevated expression of MMP-11 in 4-OHT-resistant cells, a finding corroborated by our *in vitro* long-term treatment model. MMP-9 was also upregulated in the 4-OHT-resistant cells, consistent with its well-implicated role in tumor progression, metastasis, and poor prognosis [8]. However, both MMP-16 and MMP-17 were downregulated in the resistant cells. MMP-17 is highly expressed in breast cancer and promotes tumor growth by aiding in EGFR activation post ligand stimulation, and inactivation of retinoblastoma [33]. The aberrant expression of these MMPs suggests a more complex role in 4-OHT resistance, where the upregulation of MMP-11 and MMP-9 may contribute to tumor aggressiveness. In contrast to the inverse correlation between ERα and MMP-11 observed in control MCF-7 cells, the 4-OHT-resistant breast cancer cells exhibited increased ER_α_ and MMP-11 expression. The shift in ER_α_ and MMP-11 expression may result from epigenetic or metabolic reprogramming associated with resistance. Alternatively, the co-expression of ER_α_ and MMP-11 could arise from the selection of a resistant cell subpopulation under selective pressure. Further investigation is needed to understand the mechanisms behind this shift. While our *in vitro* findings provide novel insights into the estrogen-mediated regulation of MMP-11 and its potential role in endocrine resistance, *in vivo* studies are necessary to establish the physiological relevance of estrogen’s suppression of MMP-11 in tumor progression. Given the prognostic significance of MMP-11 expression in cancer-associated stromal cells and its nuanced role in cancer progression, *in vivo* models will enable us to determine how estrogen regulation of MMP-11 influences the TME and affects cancer progression and metastasis.

In conclusion, our work demonstrates the estrogen regulation of MMP-11 expression, and suggests a protective role of estrogen in preventing cancer invasiveness. These appear to be cell-type dependent. ERα depletion results in MMP-11 upregulation, potentially driving the transition to a more aggressive phenotype. This raises important questions about whether endocrine treatments, while targeting ERα-positive tumors, might inadvertently enhance aggressive behavior by inducing MMP-11. Furthermore, 4-OHT resistance is associated with altered regulation of MMP-11, suggesting a more complex interplay of mechanisms involved in resistance and metastasis.

## Acknowledgments

The authors thank Indian Council of Medical Research, Govt. Of India, and Department of Science and Technology, Govt. Of India for financial support. Infrastructural support from Department of Biosciences and Bioengineering, IIT Guwahati is acknowledged.

## References

[1] X. Zhang, S. Huang, J. Guo, L. Zhou, L. You, T. Zhang, Y. Zhao, Insights into the distinct roles of MMP-11 in tumor biology and future therapeutics (Review), Int. J. Oncol. 48 (2016) 1783–1793. 10.3892/ijo.2016.3400.

[2] K.W. Min, D.H. Kim, S.I. Do, J.S. Pyo, K. Kim, S.W. Chae, J.H. Sohn, Y.H. Oh, H.J. Kim, S. Hyeong Choi, Y.J. Choi, C.H. Park, Diagnostic and prognostic relevance of mmp-11 expression in the stromal fibroblast-like cells adjacent to invasive ductal carcinoma of the breast, Ann. Surg. Oncol. 20 (2013). 10.1245/s10434-012-2734-3.

[3] L. Nakopoulou, E.G. Panayotopoulou, I. Giannopoulou, P. Alexandrou, S. Katsarou, P. Athanassiadou, A. Keramopoulos, Stromelysin-3 protein expression in invasive breast cancer: Relation to proliferation, cell survival and patients’ outcome, Mod. Pathol. 15 (2002) 1154–1161. 10.1097/01.MP.0000037317.84782.CD.

[4] C.W. Cheng, J.C. Yu, H.W. Wang, C.S. Huang, J.C. Shieh, Y.P. Fu, C.W. Chang, P.E. Wu, C.Y. Shen, The clinical implications of MMP-11 and CK-20 expression in human breast cancer, Clin. Chim. Acta 411 (2010) 234–241. 10.1016/j.cca.2009.11.009.

[5] Y. Zhuang, X. Li, P. Zhan, G. Pi, G. Wen, MMP11 promotes the proliferation and progression of breast cancer through stabilizing Smad2 protein, Oncol. Rep. 45 (2021). 10.3892/or.2021.7967.

[6] S.U. Kang, S.Y. Cho, H. Jeong, J. Han, H.Y. Chae, H. Yang, C.O. Sung, Y. La Choi, Y.K. Shin, M.J. Kwon, Matrix metalloproteinase 11 (MMP11) in macrophages promotes the migration of HER2-positive breast cancer cells and monocyte recruitment through CCL2–CCR2 signaling, Lab. Investig. 102 (2022) 376–390. 10.1038/s41374-021-00699-y.

[7] Y.J. Kwon, D.R. Hurst, A.D. Steg, K. Yuan, K.S. Vaidya, D.R. Welch, A.R. Frost, Gli1 enhances migration and invasion via up-regulation of MMP-11 and promotes metastasis in ERα negative breast cancer cell lines, Clin. Exp. Metastasis 28 (2011) 437–449. 10.1007/s10585-011-9382-z.

[8] M.J. Kwon, Matrix metalloproteinases as therapeutic targets in breast cancer., Front. Oncol. 12 (2022) 1108695. 10.3389/fonc.2022.1108695.

[9] N. Eiró, B. Fernandez-Garcia, J. Vázquez, J.M. Del Casar, L.O. González, F.J. Vizoso, A phenotype from tumor stroma based on the expression of metalloproteases and their inhibitors, associated with prognosis in breast cancer., Oncoimmunology 4 (2015) e992222. 10.4161/2162402X.2014.992222.

[10] A.C. Noël, O. Lefebvre, E. Maquoi, L. VanHoorde, M.P. Chenard, M. Mareel, J.M. Foidart, P. Basset, M.C. Rio, Stromelysin-3 expression promotes tumor take in nude mice, J. Clin. Invest. 97 (1996) 1924–1930. 10.1172/JCI118624.

[11] K.L. Andarawewa, A. Boulay, R. Masson, C. Mathelin, I. Stoll, C. Tomasetto, M.-P. Chenard, M. Gintz, J.-P. Bellocq, M.-C. Rio, Dual stromelysin-3 function during natural mouse mammary tumor virus-ras tumor progression., Cancer Res. 63 (2003) 5844–9. http://www.ncbi.nlm.nih.gov/pubmed/14522908.

[12] C. Su, W. Wang, C. Wang, IGF-1-induced MMP-11 expression promotes the proliferation and invasion of gastric cancer cells through the JAK1/STAT3 signaling pathway., Oncol. Lett. 15 (2018) 7000–7006. 10.3892/ol.2018.8234.

[13] E. Buache, R. Thai, C. Wendling, F. Alpy, A. Page, M.-P. Chenard, V. Dive, M. Ruff, A. Dejaegere, C. Tomasetto, M.-C. Rio, Functional relationship between matrix metalloproteinase-11 and matrix metalloproteinase-14., Cancer Med. 3 (2014) 1197–210. 10.1002/cam4.290.

[14] T. Eiseler, H. Döppler, I.K. Yan, S. Goodison, P. Storz, Protein kinase D1 regulates matrix metalloproteinase expression and inhibits breast cancer cell invasion, Breast Cancer Res. 11 (2009) 1–12. 10.1186/bcr2232.

[15] G. Kasper, M. Reule, M. Tschirschmann, N. Dankert, K. Stout-Weider, R. Lauster, E. Schrock, D. Mennerich, G.N. Duda, K.E. Lehmann, Stromelysin-3 over-expression enhances tumourigenesis in MCF-7 and MDA-MB-231 breast cancer cell lines: Involvement of the IGF-1 signalling pathway, BMC Cancer 7 (2007) 1–12. 10.1186/1471-2407-7-12.

[16] S. Nair, G. Sachdeva, Estrogen matters in metastasis, Steroids 138 (2018) 108–116. 10.1016/j.steroids.2018.07.006.

[17] Z. Piperigkou, N.K. Karamanos, Estrogen receptor-mediated targeting of the extracellular matrix network in cancer, Semin. Cancer Biol. 62 (2020) 116–124. 10.1016/j.semcancer.2019.07.006.

[18] P. Bouris, S.S. Skandalis, Z. Piperigkou, N. Afratis, K. Karamanou, A.J. Aletras, A. Moustakas, A.D. Theocharis, N.K. Karamanos, Estrogen receptor alpha mediates epithelial to mesenchymal transition, expression of specific matrix effectors and functional properties of breast cancer cells, Matrix Biol. 43 (2015) 42–60. 10.1016/j.matbio.2015.02.008.

[19] O.C. Kousidou, A. Berdiaki, D. Kletsas, A. Zafiropoulos, A.D. Theocharis, G.N. Tzanakakis, N.K. Karamanos, Estradiol-estrogen receptor: A key interplay of the expression of syndecan-2 and metalloproteinase-9 in breast cancer cells, Mol. Oncol. 2 (2008) 223–232. 10.1016/j.molonc.2008.06.002.

[20] P. Chomczynski, N. Sacchi, The single-step method of RNA isolation by acid guanidinium thiocyanate-phenol-chloroform extraction: twenty-something years on., Nat. Protoc. 1 (2006) 581–5. 10.1038/nprot.2006.83.

[21] K.J. Livak, T.D. Schmittgen, Analysis of relative gene expression data using real-time quantitative PCR and the 2-ΔΔCT method, Methods 25 (2001) 402–408. 10.1006/meth.2001.1262.

[22] N. Likhite, U.M. Warawdekar, A unique method for isolation and solubilization of proteins after extraction of RNA from tumor tissue using trizol., J. Biomol. Tech. 22 (2011) 37–44. http://www.ncbi.nlm.nih.gov/pubmed/21455480.

[23] O.H. Lowry, N.J. Rosebrough, A.L. Farr, R.J. Randall, Protein measurement with the Folin phenol reagent., J. Biol. Chem. 193 (1951) 265–75. http://www.ncbi.nlm.nih.gov/pubmed/14907713.

[24] D.J.S. John Mary, G. Sikarwar, A. Kumar, A.M. Limaye, Interplay of ERα binding and DNA methylation in the intron-2 determines the expression and estrogen regulation of cystatin A in breast cancer cells., Mol. Cell. Endocrinol. 504 (2020) 110701. 10.1016/j.mce.2020.110701.

[25] E. Afgan, D. Baker, B. Batut, M. Van Den Beek, D. Bouvier, M. Ech, J. Chilton, D. Clements, N. Coraor, B.A. Grüning, A. Guerler, J. Hillman-Jackson, S. Hiltemann, V. Jalili, H. Rasche, N. Soranzo, J. Goecks, J. Taylor, A. Nekrutenko, D. Blankenberg, The Galaxy platform for accessible, reproducible and collaborative biomedical analyses: 2018 update, Nucleic Acids Res. 46 (2018) W537–W544. 10.1093/nar/gky379.

[26] C.J. Jones, M. Subramaniam, M.J. Emch, E.S. Bruinsma, J.N. Ingle, M.P. Goetz, J.R. Hawse, Development and Characterization of Novel Endoxifen-Resistant Breast Cancer Cell Lines Highlight Numerous Differences from Tamoxifen-Resistant Models., Mol. Cancer Res. 19 (2021) 1026–1039. 10.1158/1541-7786.MCR-20-0872.

[27] G. Bhatt, A. Gupta, L. Rangan, A. Mukund Limaye, Global transcriptome analysis reveals partial estrogen-like effects of karanjin in MCF-7 breast cancer cells, Gene 830 (2022) 146507. 10.1016/j.gene.2022.146507.

[28] J.A. Aka, S.-X. Lin, Comparison of functional proteomic analyses of human breast cancer cell lines T47D and MCF7., PLoS One 7 (2012) e31532. 10.1371/journal.pone.0031532.

[29] H.S. Kim, M.G. Kim, K.W. Min, U.S. Jung, D.H. Kim, High MMP-11 expression associated with low CD8+ T cells decreases the survival rate in patients with breast cancer, PLoS One 16 (2021) 1–16. 10.1371/journal.pone.0252052.

[30] A. Ahmad, A. Hanby, E. Dublin, R. Poulsom, P. Smith, D. Barnes, R. Rubens, P. Anglard, I. Hart, Stromelysin 3:An independent prognostic factor for relapse-free survival in node-positive breast cancer and demonstration of novel breast carcinoma cell expression, Am. J. Pathol. 152 (1998) 721–728.

[31] M. Maynadier, P. Nirdé, J.-M. Ramirez, A.M. Cathiard, N. Platet, M. Chambon, M. Garcia, Role of estrogens and their receptors in adhesion and invasiveness of breast cancer cells., Adv. Exp. Med. Biol. 617 (2008) 485–91. 10.1007/978-0-387-69080-3_48.

[32] F. Weiss, D. Lauffenburger, P. Friedl, Towards targeting of shared mechanisms of cancer metastasis and therapy resistance., Nat. Rev. Cancer 22 (2022) 157–173. 10.1038/s41568-021-00427-0.

[33] E. Muñoz-Sáez, N. Moracho, A.I.R. Learte, A. Collignon, A.G. Arroyo, A. Noel, N.E. Sounni, C. Sánchez-Camacho, Molecular Mechanisms Driven by MT4-MMP in Cancer Progression., Int. J. Mol. Sci. 24 (2023). 10.3390/ijms24129944.

