## Supplementary data 1 for "Expression of MMP-11, an estrogen-suppressed gene in MCF-7 cells, is elevated upon acquisition of tamoxifen resistance"

### MMP-11 cloning, protein expression, antibody generation, and validation.

Total RNA extracted from MCF-7 cells was used for cDNA synthesis, as described in Section 2.4 (main article). The cDNA served as the template to amplify the active MMP-11 cDNA (residues 98-488, NP_005931.2) using the following primers- Forward: 5'- GGAATTCCATATGATGTTCGTGCTTTCTGGCGGGC-3', and Reverse: 5'- CCGCTCGAGTCAGAGGAAAGTGTTGGCAGG-3'. The amplified product was cloned into the pET-28a (+) vector between the NdeI and XhoI restriction sites (Fig. S1). DNA sequencing confirmed the successful insertion of the sequence.

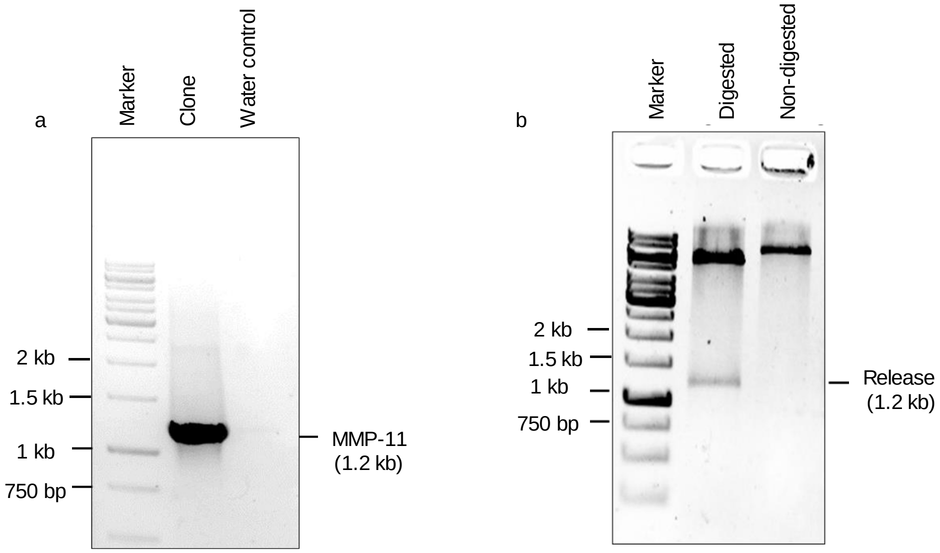


**Fig. S1. Confirmation of MMP-11 cloning into pET-28a (+) by PCR and restriction digestion. a**. Confirmation of insert in pET-28 a (+) with PCR using MMP-11 cloning primers. **b**. Restriction digestion of the clone with NdeI and XhoI showed a 1.2 kb release, confirming successful cloning.

The plasmid was then sent to BioBharati LifeScience, India, for the subsequent steps in the generation of a polyclonal MMP-11 antibody. Recombinant MMP-11 was expressed in BL21(DE3) Rosetta cells, and purified using Ni-NTA agarose beads (Fig. S2a). Rabbits were immunized with the purified recombinant MMP-11 protein. After five booster immunizations, sera were collected, tested, and processed for affinity purification. The purified antibody titer was determined using ELISA (Fig. S2b).

a b

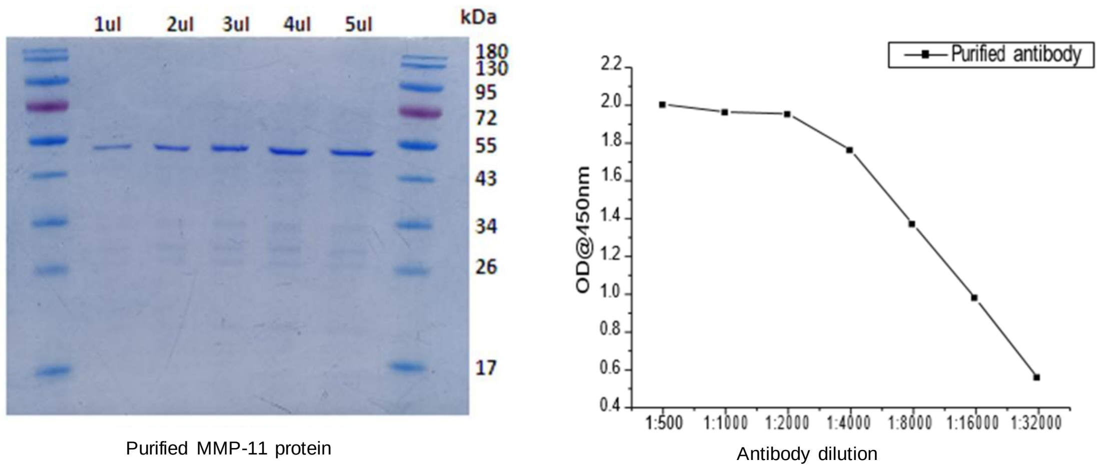


**Fig. S2. Purification of Recombinant MMP-11 Protein and Antibody Titer Determination. a.** Purified recombinant MMP-11 protein was subjected to SDS-PAGE, which showed the expected molecular weight for MMP-11. **b**. The titer of the purified antibody raised against MMP-11 was determined using ELISA. Serial dilutions of the antibody were tested against a fixed concentration of the antigen, and the absorbance was measured

Specificity was confirmed by western blotting (Fig. S3b), which showed the expected 46 kDa MMP-11 band. Western blots of total protein from MCF-7 cells treated with MMP-11 specific siRNA (Eurogentec, Belgium), showed a reduction in the 60 kDa band corresponding to the pro-enzyme form of MMP-11, further validating the antibody's specificity (Fig. S4).

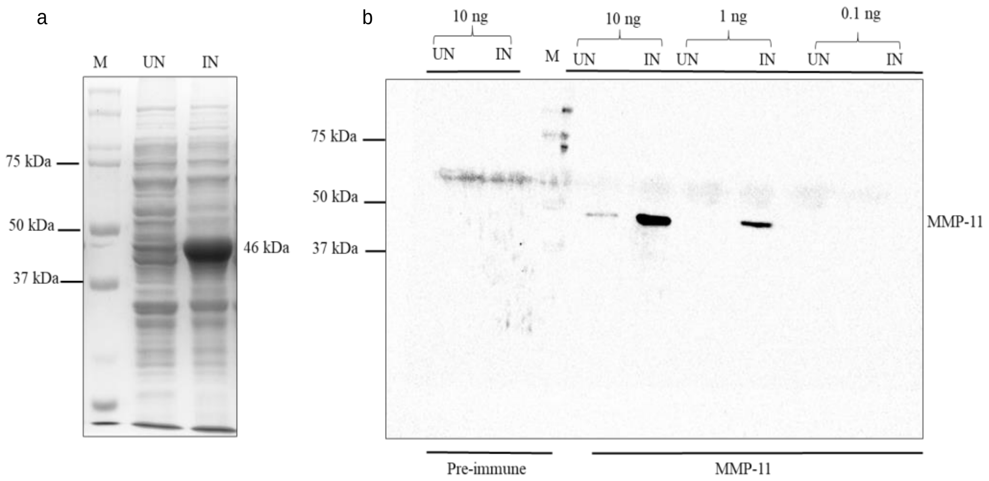


**Fig. S3. IPTG-induced expression of MMP-11 in bacterial cell lysates and detection by western blotting. a**. Coomassie-stained gel of bacterial cell lysates. IPTG induction (IN) resulted in MMP-11 expression (46 kDa). **b**. Western blotting with different concentrations (0.1-10 ng) of bacterial cell lysates and probed with MMP-11 specific antibody. The antibody detects up to 1 ng of the protein. The pre-immune sera showed no specific bands. UN indicates un-induced cell lysates and IN indicates IPTG-induced cell lysates

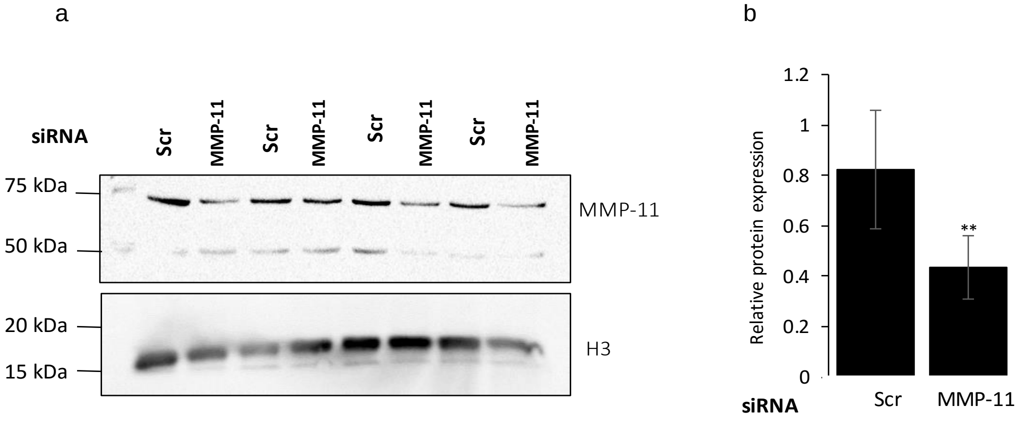


**Fig. S4. Verification of MMP-11 antibody specificity by MMP-11 knockdown**. **a,b.** Western blotting post-MMP-11 knockdown. MCF-7 cells were transfected with scrambled (Scr) or MMP-11 specific siRNA (Eurogentac, Belgium) and lysed after 96 h using RIPA lysis buffer. 30 µg of each protein sample was subjected to western blot using MMP-11-specific antibody (a). Chemiluminescence signals were processed and quantified using ImageJ software. Histone H3 served as an internal control. Quantitative representation (b) of MMP-11 protein expression. For each sample, the background-subtracted integrated band intensity for MMP-11 was normalized against that obtained for Histone (H3). The normalized MMP-11 expression in a control sample was set to one, and MMP-11 expression in other samples was expressed relative to this control. Bars represent mean relative expression ± sd (n=4). The data was analyzed by Welch's two-sample *t*-test. ** denotes significant result ($p$<0.01) relative to the control.
