## Supplementary figure 1 for "Expression of MMP-11, an estrogen-suppressed gene in MCF-7 cells, is elevated upon acquisition of tamoxifen resistance"

**Estrogen regulation of MMP-11 in T47D cells**

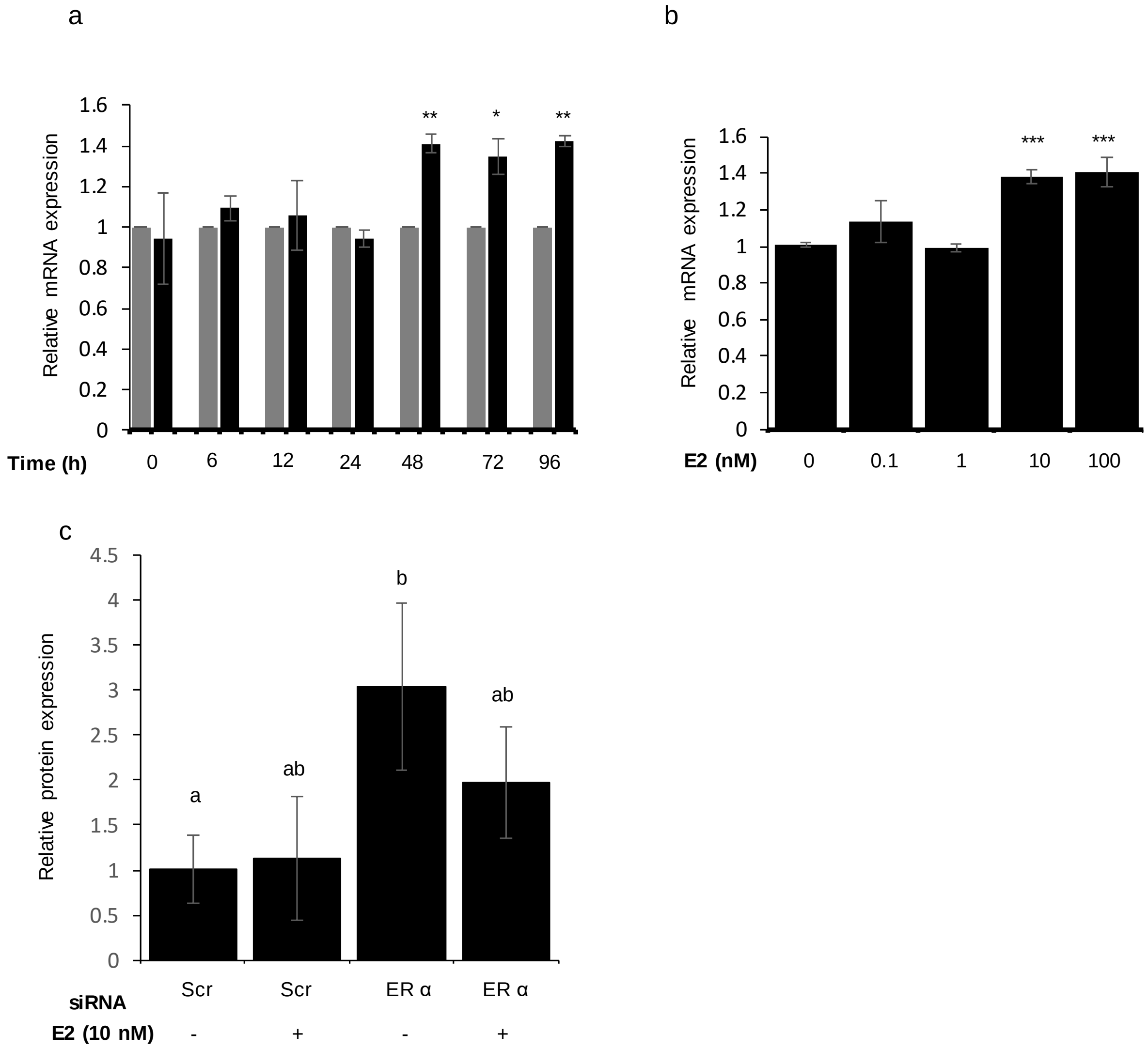


**Supplementary figure 1. E2 regulates MMP-11**. **a**. Time course experiment. T47D cells were treated with 10 nM E2 or vehicle (0.1 % EtOH) for the indicated period of time. Total RNA was isolated and subjected to RT-qPCR using MMP-11-specific primers (a). RPL35a served as an internal control. For each time point, the normalized MMP-11 expression in control samples was set to 1, and those in treated samples were expressed relative to control. Bars represent mean relative expression ± sd (n=3 biological replicates). The data for each time point were analyzed using Welch's two-sample *t*-test. Asterisks (*) and (**) denotes significant result relative to control ($p<0.05$ and $p<0.01$, respectively). **b.** Dose-response experiments. T47D cells were treated with vehicle (0.1 % EtOH) or indicated concentrations of E2 for 72 h. Total RNA was isolated and subjected to RT-qPCR using MMP-11-specific primers (b). RPL35a served as an internal control. The expression level of a control sample was set to one, and the MMP-11 expression of other samples was expressed relative to this control. Bars represent mean relative expression ± sd (n=3 biological replicates). The data were analyzed using ANOVA followed by Tukey's HSD. ***denotes significant results relative to control ($p<0.001$). **c.** Effect of ERα knockdown on E2 regulation of MMP-11. T47D cells were transfected with scrambled (Scr) or ERα specific siRNA and incubated for 24 h, followed by treatment with vehicle (0.1 % EtOH) or 10 nM E2 for 72 h. Total protein was extracted, and 30 µg of each protein sample was subjected to western blot using MMP-11 antibody. Chemiluminescence signals were processed and quantified using ImageJ software. Quantitative representation (c) of MMP-11 protein expression. For each sample, the background-subtracted integrated band intensity for MMP-11 was normalized against that obtained for Histone (H3). The normalized MMP-11 expression in a control sample was set to one, and MMP-11 expression in other samples was expressed relative to this control. Bars represent mean relative expression ± sd (n=3 biological replicates). The data were analyzed using two-way ANOVA to determine the main effects of estrogen, ERα siRNA, or their interaction, followed by Tukey's HSD post-hoc test. Letter codes (a, b, c, and d) above the bars denote statistical differences between treatment pairs
