## Supplementary Table S1 for "Expression of MMP-11, an estrogen-suppressed gene in MCF-7 cells, is elevated upon acquisition of tamoxifen resistance"

**Supplementary Table S1. List of primers**

| Sl. No | Gene Name | Primer sequence (5´→3´) | Amplicon (base pair) | Annealing temperature (°C) | Purpose |
|---|---|---|---|---|---|
| **1** | ***RPL35a*** | Forward- CGGCCTCCAAGCTCTCTAAG<br>Reverse- CAGGTCCAGGGGCTTGTACT | 131 | 60 | qRT-PCR |
| **2** | ***MMP-11*** | Forward- CCCCAAGACTCACCGAGAAG<br>Reverse- AGGTCTGTGCCCTGGTCATC | 81 | 60 | qRT-PCR |
| **3** | ***MMP11_ERE*** | Forward- CAAGTCAGTGTCACCCCCAG<br>Reverse- CCTGTCCCTGTCATGTGAGC | 140 | 60 | ChIP |
| **4** | ***pS2_ERE*** | Forward- CATTGCCTCCTCTCTGCTCC<br>Reverse- ACTGTTGTCACGGCCAAGCC | 423 | 60 | ChIP positive control |
